# Computational identification of human biological processes and protein sequence motifs putatively targeted by SARS-CoV-2 proteins using protein-protein interaction networks

**DOI:** 10.1101/2020.09.29.318931

**Authors:** Rachel Nadeau, Soroush Shahryari Fard, Amit Scheer, Emily Roth, Dallas Nygard, Iryna Abramchuk, Yun-En Chung, Steffany A. L. Bennett, Mathieu Lavallée-Adam

**Affiliations:** Department of Biochemistry, Microbiology and Immunology and Ottawa Institute of Systems Biology, Faculty of Medicine, University of Ottawa, 451 Smyth Road, Ottawa, Ontario, K1H 8M5, Canada; Department of Chemistry and Biomolecular Sciences, Centre for Catalysis and Research Innovation, University of Ottawa, Ottawa, ON, Canada

**Keywords:** COVID-19, SARS-CoV-2, Protein-Protein Interaction Network, Motif Discovery, Gene Ontology, Enrichment Analysis, Clustering, Statistics, Graph Theory

## Abstract

While the COVID-19 pandemic is causing important loss of life, knowledge of the effects of the causative SARS-CoV-2 virus on human cells is currently limited. Investigating protein-protein interactions (PPIs) between viral and host proteins can provide a better understanding of the mechanisms exploited by the virus and enable the identification of potential drug targets. We therefore performed an in-depth computational analysis of the interactome of SARS-CoV-2 and human proteins in infected HEK293 cells published by Gordon et al. to reveal processes that are potentially affected by the virus and putative protein binding sites. Specifically, we performed a set of network-based functional and sequence motif enrichment analyses on SARS-CoV-2-interacting human proteins and on a PPI network generated by supplementing viral-host PPIs with known interactions. Using a novel implementation of our GoNet algorithm, we identified 329 Gene Ontology terms for which the SARS-CoV-2-interacting human proteins are significantly clustered in the network. Furthermore, we present a novel protein sequence motif discovery approach, LESMoN-Pro, that identified 9 amino acid motifs for which the associated proteins are clustered in the network. Together, these results provide insights into the processes and sequence motifs that are putatively implicated in SARS-CoV-2 infection and could lead to potential therapeutic targets.

## INTRODUCTION

The COVID-19 (Coronavirus Disease 2019) pandemic is causing massive loss of life around the globe and has had a dramatic impact on the healthcare systems and economies worldwide. The virus at the root of this pandemic, SARS-CoV-2, is a highly pathogenic coronavirus^1^ that spreads with great efficiency^2^. While vaccines are currently in development to contain the pandemic, the efficacy of those vaccines is still undetermined^3^ and their worldwide availability will take time. Furthermore, while the therapeutic agent Remdesivir has been shown to the reduce recovery time of hospitalized COVID-19 patients, no drugs are currently approved for treatment^4^. Adding efficacious drugs to our arsenal to fight SARS-CoV-2 infections would strengthen our ability to dampen the impacts of the pandemic. Insights into the host processes that are targeted during SARS-CoV-2 infection would improve our drug development capabilities and potentially suggest drugs, for which safety has already been determined, that could be repurposed for use against SARS-CoV-2. It is therefore critical to derive a better understanding of the mechanisms by which SARS-CoV-2 infects and causes disease in human host cells. Furthermore, SARS-CoV-2 constitutes the third highly pathogenic coronavirus, which has presented a serious threat to the globe, SARS-CoV and MERS-CoV also having caused significant loss of life in the past^5,6^. Nevertheless, very little is known about their respective mechanisms, nor do drugs exist to treat their infections. The recurrence of pathogenic coronavirus outbreaks suggests a significant likelihood of another pathogenic coronavirus emerging in the future. Hence, any understanding we would gain on the SARS-CoV-2 infection and identification of compounds that are effective for use as a treatment could play a critical role in containing future pathogenic coronavirus outbreaks.

Protein-protein interactions (PPIs) are extremely useful to map out biological processes, protein machineries, and protein complexes^7^. In an effort to provide a better understanding of the biological processes affected by SARS-CoV-2 in human host cells, Gordon et al. mapped the interactions of SARS-CoV-2 proteins with human proteins in infected HEK 293 cells. Their effort early on during the outbreak highlighted several potential drug targets and drug candidates. It also generated an interactome containing a great wealth of information that can be further analyzed to identify novel insights into SARS-CoV-2 mechanisms of infection and host-pathogen interactions.

Algorithms including the Markov Clustering algorithm (MCL)^8^, Restricted Neighborhood Search Clustering Algorithm (RNSC)^7^, MCODE^9^, Socio-Affinity Index^10^, and the eigenmode analysis of the connectivity matrix of a PPI network^11^ have proven effective at characterizing protein complexes in PPI networks using clustering approaches. Such clusters can then be investigated for functional enrichment by determining whether a given function is overrepresented among the proteins within the cluster with respect to the number of proteins it annotates in the entire network. Gene Ontology terms^12^, KEGG pathways^13^, and REACTOME pathways^14^ are often used for such enrichment analyses.

We have shown in the past with our tool GoNet that evaluating the clustering of GO terms in protein-protein interaction networks can be done to identify biological processes and protein complexes of interest^15^. We have also shown that PPI networks can be effectively used to discover novel RNA sequence motifs that are associated with groups of significantly clustered proteins using another algorithm called LESMoN^16^. Herein, we propose a set of functional enrichment analyses and protein sequence motif discovery approaches to thoroughly investigate the set of human proteins interacting with SARS-CoV-2 proteins revealed by Gordon et al. We first directly analyze their network of PPIs to identify GO terms and protein sequence motifs that are enriched amongst SARS-CoV-2-interacting proteins. We then supplement this set of interactions with known human PPIs and apply clustering approaches to identify GO terms and protein sequence motifs that are clustered in the network. Given the relevance of protein-protein interactions in defining the effects of viral proteins on host processes, these functional annotations and sequence motifs are likely to reveal previously unappreciated aspects of SARS-CoV-2 infection. Our analyses provide a better understanding about the effect of SARS-CoV-2 on host cell machinery and have the potential to help in the discovery or repurposing of drugs targeting COVID-19.

## EXPERIMENTAL SECTION

### Overview

We present a suite of enrichment analyses investigating both functional and sequence motif overrepresentation (Figure 1). We directly analyze the viral-host PPI network generated by Gordon et al. and also build a second network from it of human PPIs using STRING. We perform GO and sequence motif enrichment analyses on the viral-host PPI network. We then identify GO terms and sequence motifs that are clustered in the human PPI network derived with the help of STRING using two clustering approaches.

**Figure 1.**
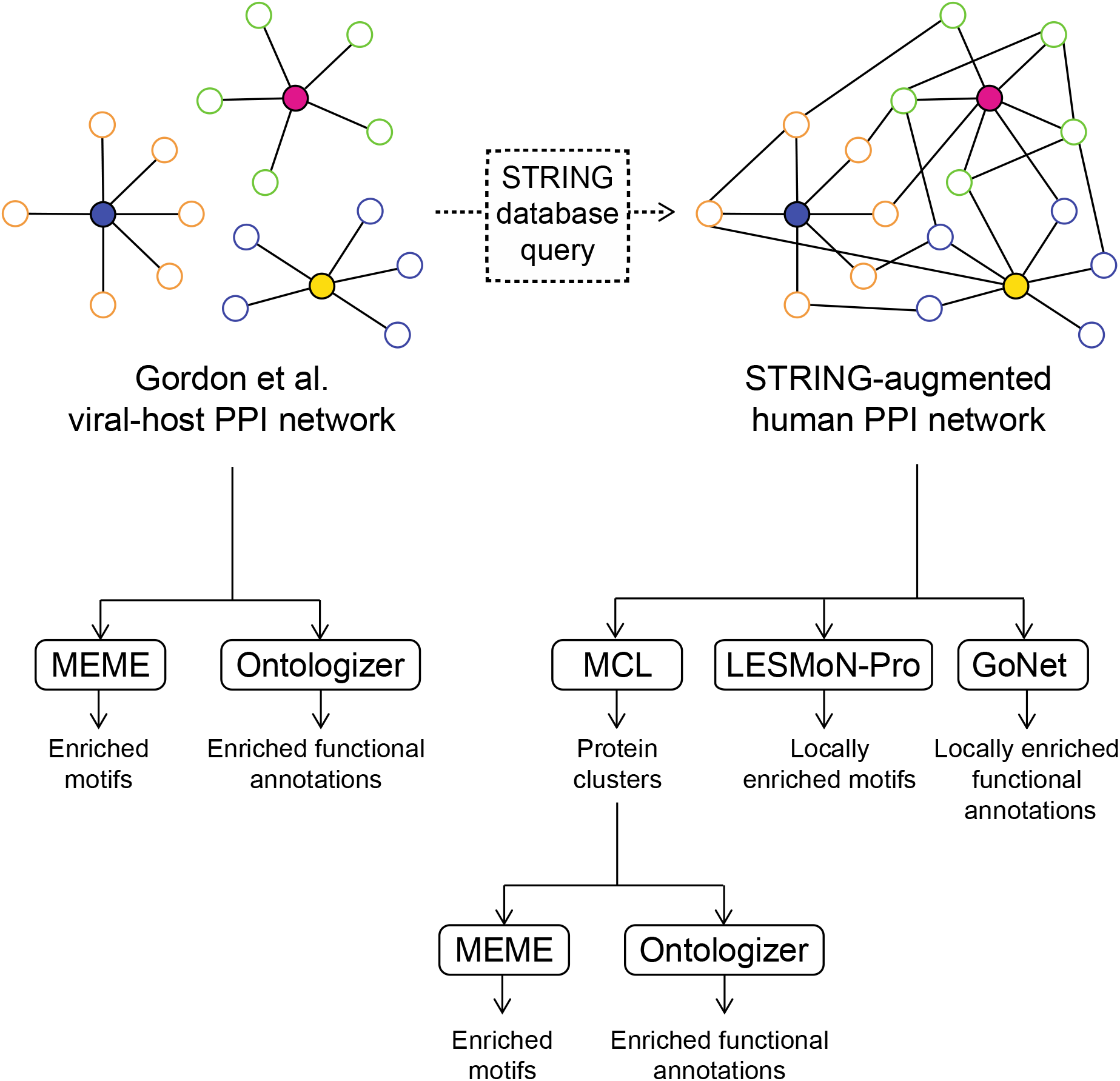
Graphical representations of the PPI networks analyzed and the enrichment analysis approaches applied on them. Color-filled proteins in the PPI networks represent SARS-CoV-2 proteins while proteins without color filling represent human proteins.

### Dataset

Our computational analyses were performed on two PPI networks. The first one is the high-confidence viral-host PPI network of SARS-CoV-2 proteins interacting with *H. sapiens* HEK293 proteins generated by Gordon et al.^17^. Since this network was built solely by the affinity purification of SARS-CoV-2 proteins, the interactions between *H. sapiens* proteins are not included in this network. We therefore built a second version of this network by adding interactions between human proteins in the viral-host PPI network based on the information stored in the STRING PPI database^18^. Human proteins that were purified by SARS-CoV-2 proteins were therefore connected if they reached a medium confidence (0.4) according to STRING and had either an experimental or database evidence. This resulted in the creation of a network containing 216 proteins and 502 PPIs (Supplementary Table S1). However, not all of these proteins are connected to each other. We therefore identified the largest connected component of the network and discarded other proteins, which resulted in a network of 195 proteins and 489 PPIs. This network will be referred to as the STRING-augmented network.

### Markov Clustering algorithm to identify clusters in the STRING-augmented network

We used the Markov Clustering algorithm (MCL)^8^ to identify clusters of proteins in the STRING-augmented network. The algorithm was executed using an inflation parameter of 2 to obtain a reasonable level of granularity in the protein clusters.

### Gene Ontology enrichment analysis

In order to identify overrepresented biological processes, cellular components, and molecular functions, a GO enrichment analysis was performed using the Ontologizer package^19^ on both the viral-host PPI network and the clusters obtained by the MCL analysis of the STRING-augmented network. An analysis was performed for each individual set of proteins purified by a SARS-CoV-2 protein and for each cluster of size 2 or greater detected by MCL. The set of all proteins present in the respective networks was used as background for the enrichment analyses. Ontologizer uses a modified Fisher’s exact test to assess the statistical significance of a GO term enrichment. The resulting *p*-values were adjusted for multiple hypothesis testing using the Benjamini-Hochberg procedure^20^.

### Protein Sequence Motif Enrichment analysis

With the objective of discovering protein sequence motifs that are surprisingly overrepresented in certain sections of the networks, we performed a motif enrichment analysis using the MEME Suite^21^ on network elements of the viral-host network and the STRING-augmented network. As for the GO enrichment analysis, MEME was executed on each individual set of proteins purified by a SARS-CoV-2 protein. It was also executed on all clusters of size 3 or more as detected by the MCL algorithm. MEME was executed in classic mode with a 0-order Markov Model and zero or one occurrences per site for motifs with a minimum width of 5 and a maximum width of 30.

### Identifying GO terms that are clustered in the STRING-augmented network using GoNet 2.0

To investigate the clustering of proteins sharing the same GO terms, we implemented a modified version of our previously published approach, named GoNet^15^.

#### Clustering significance assessment

Briefly, as previously described, GoNet measures the clustering of a GO term *t*, annotating *P* proteins by calculating its total pairwise distance (TPD) in the network, which in this context is the shortest path between all pairs of proteins annotated with *t*. GoNet then assesses the significance of this clustering measure using a Monte Carlo sampling approach. We modified GoNet’s statistical approach to take into account the annotation bias of proteins. To assess the significance of the total pairwise shortest path of *t*, GoNet randomly samples without replacement *P* proteins in the network 100,000 times and computes the total pairwise distance between all proteins in each sample. The difference in this new version of GoNet is that the sampling probability of each protein is proportional to the number of GO terms annotating them. This procedure ensures that proteins that are annotated by a large number of GO terms are sampled more often than those with fewer GO terms, since the former proteins will see their clustering tested more often in the network. Using an approach inspired by our clustering significance assessment tool, LESMoN^16^, we estimated the mean and standard deviation of the TPD of each value of *P* and derived a normal distribution of the TPD for all *P*’s. These distributions can be used as a null model to estimate a *p*-value for the TPDs of all GO terms.

#### False discovery rate estimation

Since GO terms share several protein annotations, the statistical testing of GO term clustering is not independent. Traditional multiple hypothesis testing correction strategies are therefore likely to be overly conservative in their *p*-value adjustments. Instead, again inspired by our LESMoN algorithm, we used a permutation-based strategy to estimate false discoveries. Briefly, pairs of GO-protein annotations are randomly selected and swapped, such that if protein A is annotated by GO term 1 and protein B is annotated by GO term 2, after their random selection and swapping, A will be annotated by GO 2 and B with GO 1. This annotation swapping procedure is performed 1,000 x the total number of GO-protein associations times, such that the annotation permuted network is representative of a randomly annotated network. After this annotation permutation, GoNet is then executed such that the statistical significance of the clustering of the permuted GO terms is assessed. False Discovery Rate (FDR) at a given p-value α threshold is then estimated as follows:

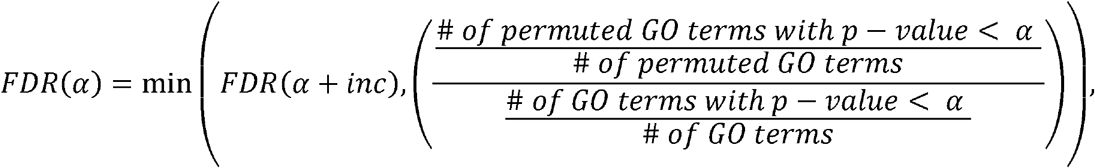

Where *inc* is a small increment in *p*-values. FDR of *α* corresponds to the minimum between the FDR estimated for a slightly higher *p*-value and the FDR at *α* such that the FDR function remains monotonic, preventing noisy FDR fluctuations at very small p-value thresholds.

### Identifying protein sequence motifs that are locally enriched in the STRING-augmented network using LESMoN-Pro

We have previously presented an approach called LESMoN, which identifies 5’ untranslated sequence motifs for which the associated proteins are clustered in a PPI network^16^. Herein, we propose a new version of LESMoN, called LESMoN-Pro, which detects protein sequence motifs for which the associated protein sequences are clustered in a PPI network. LESMoN-Pro uses similar algorithmic principles as LESMoN, while being adapted to search for protein motifs instead of RNA motifs. Briefly, LESMoN-Pro enumerates amino acid motifs of size 8 over the following alphabet: A, C, D, E, F. G, H, I, K, L, M, N, P, Q, R, S, T, V, W, Y, X, B, Z, J, l, @, h, o, p, t, s, b, +, -, c, wherein the following are degenerate characters encompassing multiple amino acids: B (asparagine and aspartic acid), Z (glutamine and glutamic acid), J (leucine and isoleucine), l (aliphatic), @ (aromatic), h (hydrophobic), o (alcohol), p (polar), t (tiny), s (small), b (bulky), + (positively charged), - (negatively charged), c (charged), and X is a wild card character corresponding to all amino acids. All other characters correspond to single amino acids. The exact sets of amino acids mapping to the different degenerate characters are given in Supplementary Table S2. Using this alphabet, LESMoN-Pro enumerates motifs that are represented in the set of protein sequences of the 195 proteins forming the largest connected component of the STRING-augmented PPI network, such that a maximum of 4 degenerate characters can appear in any given motif. This rule ensures that the motifs are not too degenerate (i.e., present in most if not all proteins) and also reduces the algorithm’s running time. Protein sequences were downloaded from the UniProt SwissProt database on Feb. 12, 2020. Motifs that matched to the sequences of at least 3 proteins were retained for downstream analyses. The clustering of the proteins associated to a total of 48,128,026 motifs was evaluated. Motifs were associated with sets of proteins ranging in size from 3 to 194 proteins.

#### Clustering significance assessment

LESMoN-Pro’s clustering significance assessment of motifs in the PPI network mirrors the one from GoNet. Indeed, one could consider that a sequence motif contained in a protein sequence is annotating that protein, much like a GO term annotates a protein. Hence, the clustering of the proteins containing a given motif was measured using the total pairwise distance given by the shortest paths between all pairs of proteins. A Monte Carlo sampling approach was then used to assess the statistical significance of the clustering. Once again, since some proteins are likely to contain more motifs than others due to their sequence length or amino acid composition, sampling of individual proteins is performed with a probability proportional to the number of motifs associated with a protein. For all sizes of sets of proteins annotated by a motif, a null distribution of the TPD is estimated using a normal distribution approximated using the mean and standard deviation obtained from 100,000 samplings of protein sets. A *p*-value is then estimated by comparing the TPD of a given motif to its corresponding null distribution.

#### False discovery rate estimation

In order to estimate an FDR at a given *p*-value threshold, protein sequences were locally shuffled as previously described^16^. Briefly, amino acids within non-overlapping sliding windows along all protein sequences were shuffled. 10,000 pairs of amino acids were randomly selected within each window and permuted. This procedure ensures that any local sequence properties are maintained while generating sequences that are not likely to be present in the human proteome. The LESMoN-Pro assessment of clustering significance is then applied on the motifs present in those locally shuffled sequences. Using the same strategy as GoNet, an FDR is estimated for a given *p*-value.

#### Determining sequence motif families of biological interest

Since a number of interacting proteins in the STRING-augmented network share important levels of sequence homology, a number of motifs found are likely to simply originate from highly homologous regions. While we acknowledge that such motifs are not devoid of biological interest, we claim that they are not as interesting, or at least surprising, as motifs conserved in interacting proteins that do not share a high level of sequence homology. We therefore built a procedure for filtering out motifs that are mainly contained in highly homologous protein sequences. Specifically, protein sequence identity between all pairs of proteins was computed by Clustal Omega^22^. Motifs for which 75% of the pairs of the associated proteins showed a sequence identity > 25% were deemed to be originating primarily from homologous sequences and discarded. Furthermore, similar motifs are likely to share a similar level of clustering significance. In order to avoid reporting redundant results, we used LESMoN’s approach of motif family grouping^16^. Specifically, a hierarchical clustering approach using average linkage and a similarity measure based on the fraction of shared proteins between pairs of motifs was used to identify families of similar motifs. A representative motif for each family is then determined using the motif with the best clustering *p*-value. In the advent of a tie, the motif that is the least substituted with degenerate characters, based on the number and specificity of the degenerate characters, is selected. If no single motif is identified as the least degenerate among the motifs with the lowest *p*-value, one motif from the list of such motifs is randomly selected as the representative motif. Finally, a consensus motif is then built using ggseqlogo^23^ based on the matching occurrences of the representative motifs in the associated protein sequences.

### Software availability

All software packages developed and implemented for GoNet and LESMoN-Pro are open-source and accessible at this address: https://github.com/LavalleeAdamLab/sars-cov-2

## RESULTS

We present an in-depth enrichment analysis of PPI networks involved in SARS-CoV-2 PPIs. We apply a GO and a motif enrichment analysis on sets of proteins that were purified by SARS-CoV-2 proteins in the Gordon et al. study. We also generate a PPI network of human proteins that are interacting with SARS-CoV-2 proteins, based on interactions derived from the STRING database. Using this network, we use clustering strategies and enrichment analysis tools to reveal functional annotations and protein sequence motifs for which the associated proteins are significantly clustered in the PPI network.

### Human proteins purified by SARS-CoV-2 proteins show functional enrichments

While Gordon et al. performed a GO enrichment analysis on the proteins they have purified using SARS-CoV-2 proteins, they only performed this analysis for biological processes. We complemented this analysis investigating both GO cellular components (Figure 2A-K) and molecular functions (Supplementary Figure S1), in addition to having repeated the analysis for biological processes using a different algorithm (i.e., Ontologizer) (Supplementary Figure S2) with the goal of providing a better understanding of the role played by SARS-CoV-2 proteins. Complete Ontologizer GO enrichment analysis results are provided in Supplementary File S1. These enrichment analyses highlighted, among other findings, that the SARS-CoV-2 protein N interactors were enriched for the term “Nucleus” (FDR-adjusted *p*-value (*p*) = 0.027), the PPIs of nsp1 were enriched for “MCM complex” proteins (*p* = 4.9 x 10^−6^), and nsp8 interactors were overrepresented by “nucleus” proteins (*p* = 0.033), while the interactors of nsp9 were enriched for “nuclear pore” proteins (*p* = 0.0014), suggesting that these proteins may play a role in the nucleus of the host cell. On the other hand, SARS-CoV-2 nsp7 show enrichment among its interactors for “membrane” (*p* = 0.0042) and “vesicle” proteins (*p* = 0.042). Interestingly, SARS-CoV-2 nsp13’s PPIs were highly enriched for microtubule organizing center proteins (*p* = 1.3 x10^−11^), while orf10’s interactors were enriched for ubiquitin ligase complex proteins (*p* = 0.0032).

**Figure 2.**
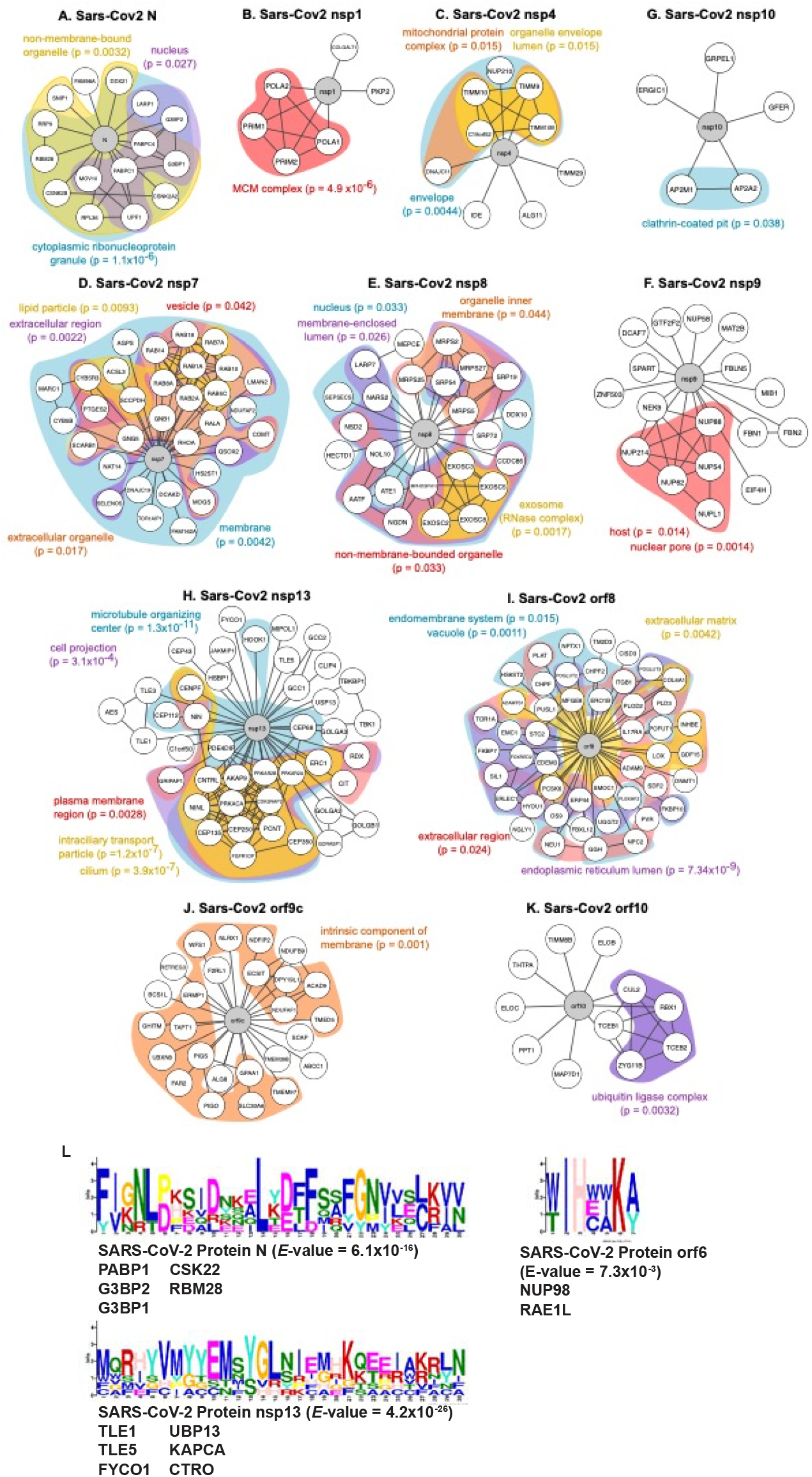
GO enrichment analysis of proteins interactors of a selected set of SARS-CoV-2 proteins. A) – K): GO cellular components are color-coded. *p* corresponds to an FDR-adjusted *p*-value (See methods). L) Motifs enriched among the protein sequence of the interactors of SARS-CoV-2 proteins N, nsp13 and orf6. Proteins containing the motifs are listed below the motif logos.

### Human proteins purified by SARS-CoV-2 proteins are enriched for specific motifs

In addition to functional enrichment, we investigated whether the groups of proteins interacting with SARS-CoV-2 proteins were enriched for specific amino acid sequence motifs using the MEME Suite^16^ (Supplementary File S2). Such motifs could play an important role in the binding of SARS-CoV-2 proteins or in the host mechanism these proteins are affecting. A number of SARS-CoV-2 proteins showed such enrichments (E-value < 0.05). Among these, SARS-CoV-2 proteins N, nsp13, and orf6 interacted with proteins sharing similar sequence motifs (Figure 2L). It is worth noting that many more motifs were identified, but a large portion of them, such as those identified for nsp7, appeared to be detected by MEME simply because the interactors of this protein, RAB-family proteins, share a high level of homology (Supplementary File S2).

### Clustering analysis reveals biological processes that are putatively affected by SARS-CoV-2

In an effort to characterize the biological processes, cellular components, and molecular functions that are affected by SARS-CoV-2 infection, we supplemented the viral-host PPI network of Gordon et al. with STRING PPIs in order to connect the human proteins based on known interactions. This enabled us to perform a clustering analysis using the MCL algorithm in order to identify proteins that are interacting with SARS-CoV-2 proteins and also closely interacting together. Our MCL analysis revealed 29 clusters of at least 3 proteins (Supplementary Table S3). A GO enrichment analysis on these clusters revealed several GO biological processes that were significantly overrepresented (FDR-adjusted *p*-value < 0.01; Figure 3A). Among the main GO biological processes enriched in these clusters, we find GO terms related to establishment of RNA localization, metabolism, membrane docking, cell cycle, among other things. We also find GO cellular components related to nuclear pores, microtubules, cilia, and cell projections to be enriched (FDR-adjusted *p*-value < 0.01; Supplementary Figure S3A). Finally, GO molecular functions such as transporter activity, lyase activity, and structural molecule activity are enriched in the clusters (FDR-adjusted *p*-value < 0.01; Supplementary Figure S4A). Complete GO enrichment analysis results of the MCL clusters are available in Supplementary File S3. These GO terms highlight the main functional annotations that are likely affected by SARS-CoV-2.

**Figure 3.**
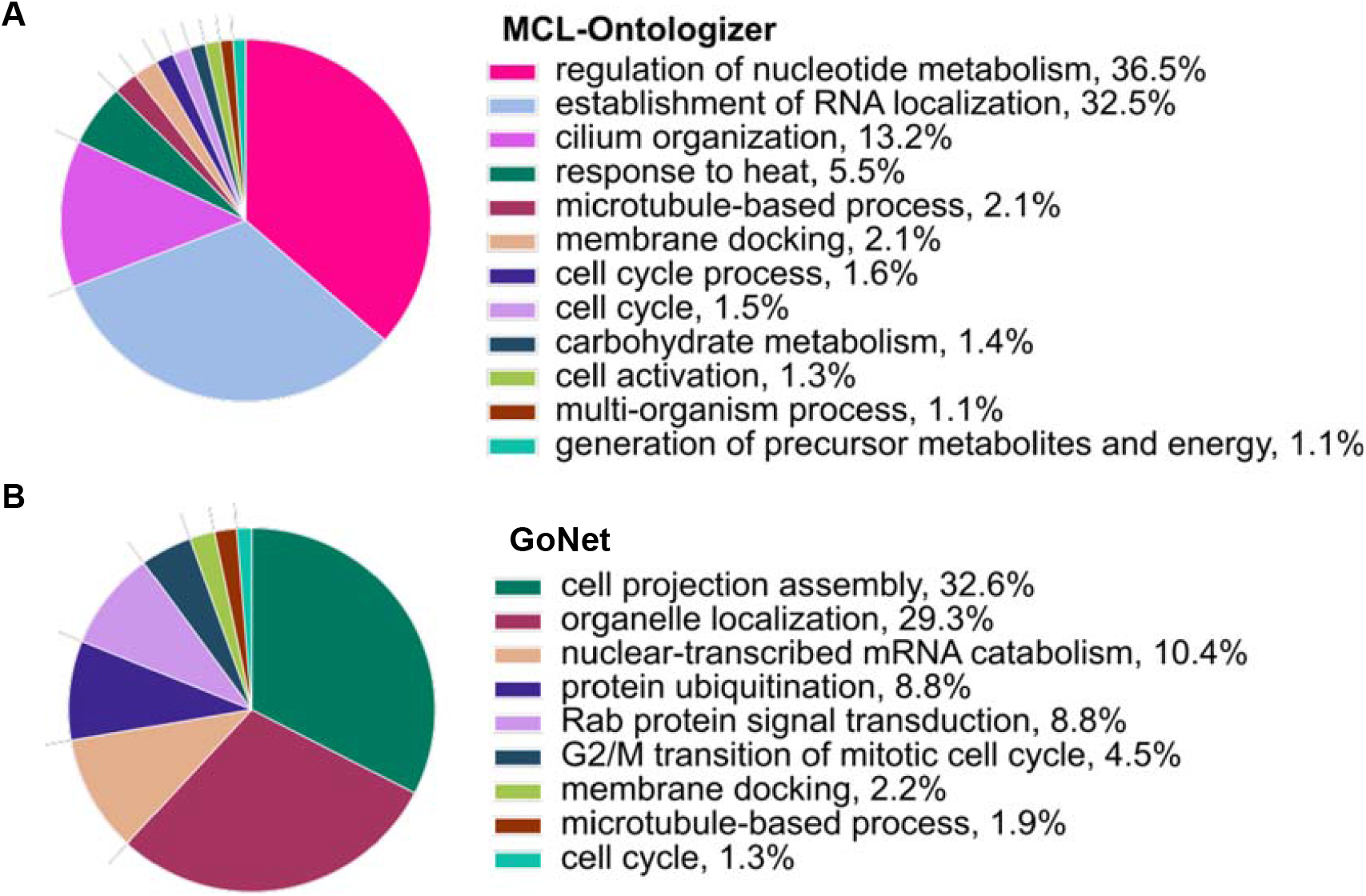
Pie charts of the GO enrichment analysis of MCL clusters (FDR-adjusted p-value < 0.01) (A) and clustering statistical significance of GO terms according to GoNet (FDR < 0.01) (B). GO biological processes with the highest level of enrichment statistical significance are attributed larger pieces of the pie. The portion occupied by each GO term is also represented in percentages next to the term names.

To further investigate the clustering of GO terms in the STRING-augmented network, we propose a novel adapted version of our GoNet algorithm, which detects GO terms for which the proteins are surprisingly clustered in a PPI network (Supplementary Table S4). The advantage of this approach over MCL is that no clusters need to be predetermined before GO enrichment analysis. Indeed, MCL could break apart a large cluster into two smaller clusters and therefore separate a given GO term enriched in the large cluster. GoNet instead simply evaluates the clustering of GO terms and is therefore not affected by overlapping clusters. Overall, GoNet identified 329 GO terms for which the associated proteins were significantly clustered in the STRING-augmented PPI network (*p*-value < 0.001 and FDR < 0.01). GoNet detected GO biological processes that MCL highlighted, but also uniquely identified GO terms related to protein ubiquitination and RAB protein signal transduction, which were found to be significantly clustered (FDR < 0.01; Figure 3B). GoNet also identified GO cellular components related to centrosomes, cell division sites, and vesicles that the MCL-based approach did not highlight (FDR < 0.01; Supplementary Figure S3B). Finally, GoNet found additional GO molecular functions related to myosin binding, protein kinase A binding, and repressing transcription factor binding to be locally enriched in the PPI network, which MCL did not highlight (FDR < 0.01; Supplementary Figure S4B). Once again, these GO term clusterings highlight groups of highly interacting proteins involved in similar functions that are known to be bound and therefore putatively affected by SARS-CoV-2.

### Protein sequence motifs are locally enriched in certain regions of the STRING-augmented PPI network

In addition to investigating GO overrepresentation in MCL-derived protein clusters, we performed a motif enrichment analysis using the MEME suite (Supplementary File S4). This analysis highlighted a number of sequence motifs that are surprisingly overrepresented within MCL clusters (E-value < 0.05; Figure 4 and Supplementary File S4). However, the overwhelming majority of these motifs are likely due to high sequence homology between proteins as can be seen from the families to which these proteins belong (i.e., RAB, TIM, TLE and CSK proteins).

**Figure 4.**
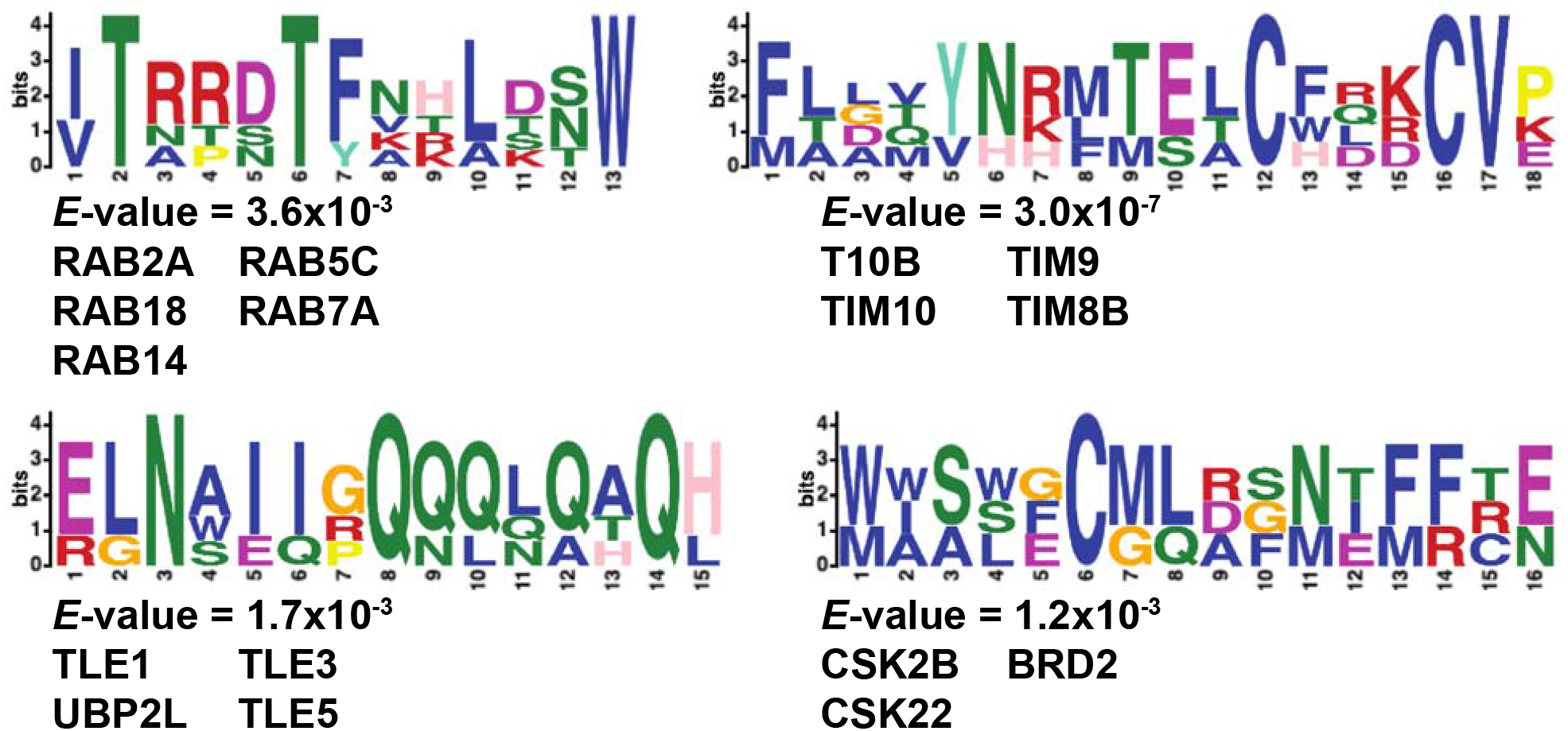
Four protein sequence motifs that were enriched in protein clusters detected by MCL in the STRING-augmented PPI network (E-value < 0.05). Proteins containing the motifs are listed below the motif logos.

### LESMoN-Pro identifies families of protein sequence motifs that are significantly clustered in the STRING-augmented network

Using the LESMoN-Pro algorithm to identify amino acid sequence motifs for which the associated proteins are significantly clustered in the STRING-augmented PPI network, we generated a list of 363,279 motifs (FDR < 0.05). After filtering this list of motifs for which fewer than 75% of the pairs of their associated proteins were homologous (see Experimental Section), 1,650 motifs remained (Supplementary Table S5). All of which contained either 3 or 4 degenerate characters. To facilitate downstream analysis of these 1,650 motifs and since a large number of motifs are likely to be very similar, we grouped them into 9 families using hierarchical clustering based on the similarity of their lists of associated proteins, and then selected a representative motif for each family (Figure 5 and Supplementary Figure S5). Figure 5A shows that the sets of annotated proteins tended to be centered around highly clustered groups of somewhat homologous proteins, including RAB, NUP, and MARK proteins, but that all included at least one additional protein that share little homology with these families. Among these proteins, both MIB1 and TBK1, which are involved in the same pathway and implicated in innate antiviral immunity, contained the same representative motif, HXXIXKLb^24^. In the SARS-CoV-2 viral-host PPI network, MIB1 and TBK1 are associated to two different viral proteins, Nsp9 and Nsp13, respectively, suggesting a potential role for the identified motif in the host response, rather than one directly affecting viral protein binding. Overall, our results identify families of sequence motifs that are significantly clustered in the STRING-augmented PPI network. Such motifs may represent viral protein binding sites and may provide insights into the host processes targeted by SARS-CoV-2 viral proteins and their interactions.

**Figure 5.**
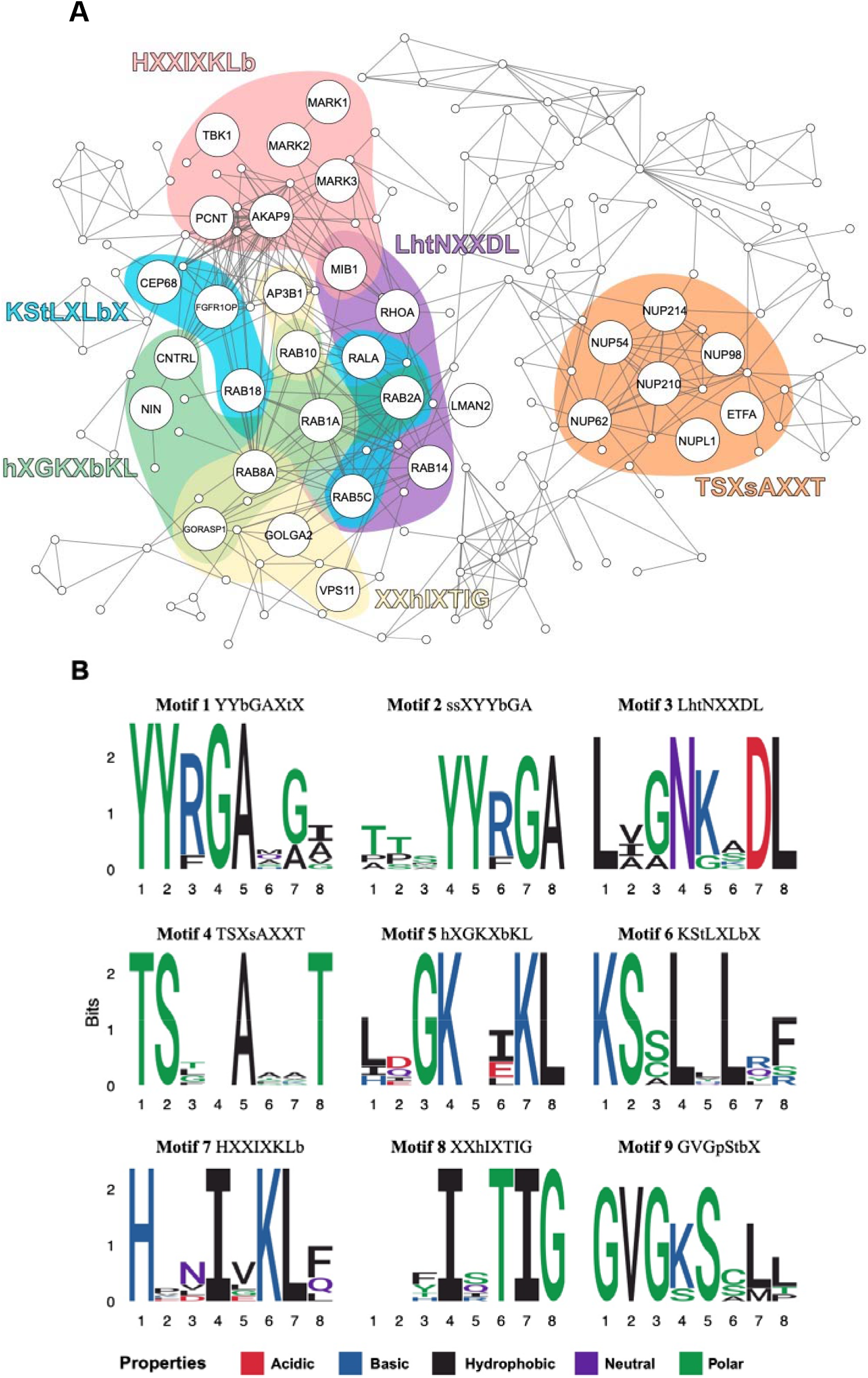
Family representative sequence motifs for which the associated proteins are significantly clustered in the STRING-augmented network. A) Complete STRING-augmented network where proteins containing significantly clustered motifs are larger and labeled. A) Selected set of representative motifs are shown on the network coloring the proteins containing them (FDR < 0.05). B) All family representative motifs detected by LESMoN-Pro are shown as sequence logos built from their actual occurrences in their associated protein sequences. Properties of amino acids are color-coded on the logos.

## DISCUSSION

### Locally enriched GO terms relevance with regards to COVID-19

Our initial analysis explored the viral-host PPI network of human proteins interacting with SARS-CoV-2 proteins. This analysis highlighted that interactors of the viral Nsp proteins were enriched for GO terms such as “MCM complex”, “Nucleus”, “Nuclear pore”. Given that Nsp proteins have been shown to play an important role in RNA replication and transcription, it is not surprising to see such associations^25,26^. Within this group of Nsp proteins, we saw a high enrichment for the “microtubule organizing center” (MTOC) GO term among Nsp13 interactors. It has been previously reported that Nsp13 is a multi-functional SARS-CoV helicase that unwinds duplex RNA/DNA and, using ATP, is able to translocate along the nucleotides^27^. It has been an early pharmaceutical target for SARS-CoV inhibition, with intention to reduce the unwinding ability of the viral helicase^28^. It was proven to be a promising molecular target for as the viral helicase could be inhibited without affecting the activity of the human helicase^29^, and this protein continues to be studied for the novel SARS-CoV-2^30,31^. However, the exact interactions between Nsp13 and MTOC have not been characterized to our knowledge. The proteins associated with this GO term may reveal further insight into the replicative mechanism of the virus and assist in the development of improved inhibitors.

We a saw a high enrichment for “ubiquitin ligase complex proteins” among the interactors of Orf10. This was also confirmed by our GoNet algorithm, which detected ubiquitination-related GO terms as significantly clustered in the network. Proteases have also been an early and promising pharmaceutical target for SARS-CoV-2 ^32–34^. The entry of the virus has been putatively linked to the ACE2 receptor and priming of the viral S protein by host proteases^35–37^. Interestingly, viral entry can be successfully blocked with the use of a protease inhibitor^33^, making this an exciting intervention option. Thus far, protease-related activity in SARS-CoV-2 has been associated with the viral S protein. Among the novel SARS-CoV-2 proteins, Orf10 is one of the least homologous to previous coronavirus proteins^38,39^ and therefore has not been as studied to the same extent. However, the association of Orf10 to protease-related functions makes this protein an interesting candidate for further investigation. Its implication in the protease pathway can give us further insight into viral protein priming by the host and can ultimately improve current pharmaceutical targets for inhibiting viral entry into the cell. These GO terms represent a few examples that are thought to be useful for therapeutic developments. However, our approach suggests many more GO terms of interest for which the associated proteins may also constitute viable drug targets. Our GO clustering analyses detected clustering of great significance for annotations such as establishment of RNA localization, cell projection, lyase activity, cell cycle and more. These fall in line with the current literature which describes such disturbances upon SARS-CoV-2 infection^40–42^ and could also represent putative therapeutic targets. This collection of GO terms also helps shed light on the host mechanisms hijacked by SARS-CoV-2.

### Locally enriched motifs relevance with regard to COVID-19

Motifs for which the associated proteins are significantly clustered in the STRING-augmented PPI network may be informative by uncovering potentially important protein sequences involved in both downstream host processes targeted by SARS-CoV-2 and in the direct interactions between viral and host proteins. An example of the former is the pair of proteins MIB1, an E3 ubiquitin-protein ligase, and TBK1, a serine-threonine kinase, both of which share the sequence motif HXXIXKLb and directly interact to regulate innate antiviral immunity by leading to the phosphorylation of interferon regulatory factors, which transcriptionally activate downstream pro-inflammatory and antiviral genes^24^. If SARS-CoV-2 suppresses antiviral immunity in host cells through interactions with these proteins, analysis of shared sequence motifs may therefore provide insights to discover other proteins that may be involved in these disrupted biological processes. Furthermore, analyzing the clustering of shared sequence motifs in the PPI network may facilitate the identification of related biological processes that are targeted by, interact with, or are affected by SARS-CoV-2 viral proteins. For example, VPS11, a protein involved in vesicle-mediated protein trafficking^43^, AP3B1, a subunit of an adaptor protein complex involved in protein sorting^44^, and GORASP1 and GOLGA2, which are involved in maintenance of^45^ and vesicle fusion^46^ to the Golgi apparatus, respectively, are significantly clustered in the PPI network and share the sequence motif XXhIXTIG. This potentially suggests a related role for all four proteins, and possibly the sequence motif itself, in the trafficking of viral components during SARS-CoV-2 infection. Finally, while the representative motif chosen for each family provides insights into some of the proteins sharing similar sequence motifs, it is important to note that it does not capture the full variety of the proteins in the network that are annotated by other motifs grouped in the same family. Therefore, further information about the sequence motifs potentially targeted by viral proteins and involved in host processes targeted by SARS-CoV-2 may be derived from further analyses of these additional motif-annotated proteins.

### The influence of highly homologous proteins in motif discovery

It is clear that some of the proteins containing the motifs discovered by MEME or LESMoN-Pro share these motifs because they are highly homologous and closely interacting with each other. While these motifs are part of larger protein sequence domains may be of biological interest to understand the mechanisms of SARS-CoV-2 infection, their occurrence is much less surprising than those in proteins that do not share a high level of homology. For the majority of the motifs reported here, while some proteins hosting these motifs share some level of homology, we observe a good fraction, at least 25% of the proteins, that do not. The fact that these proteins share a high degree of clustering and the presence of a given motif despite sharing a low level of homology make them interesting research avenues to provide a better understanding of SARS-CoV-2 infection. In the future, it could be of interest to mutate such motifs and observe their impact on SARS-CoV-2 infection.

### Reliability of PPIs

In this work, we have considered all PPIs to be of equal quality. However, some show a higher level of confidence and some occur at a greater frequency than others. While we have little information about the latter, approaches considering the former to weight PPI networks based on such reliability could be designed. Tools such as SAINT^47,48^, ComPASS^49^, and Decontaminator^50,51^ could be used to obtain reliability scores that could be transformed into weights for the interactions in the network. MCL, GoNet, and LESMoN-Pro could therefore be adapted to consider such weights when performing their clustering analysis to reveal more biologically relevant clusters. While such an analysis may be rewarding, it would however remain challenging due to the highly heterogenous nature of PPIs deposited into STRING.

## CONCLUSION

In this manuscript, we report an ensemble of GO terms (biological processes, cellular components and molecular functions) that are likely impacted by SARS-CoV-2 infection in human. These annotations and their associated proteins can help provide a better understanding of the mechanisms exploited by SARS-CoV-2. We also highlight 9 novel protein sequence motifs that are likely to be closely affected by SARS-CoV-2. Such motifs could represent binding sites or SARS-CoV-2 or play key roles in the processes hijacked by the virus.

## SUPPORTING INFORMATION

The following files are available free of charge.

**Supplementary Figure S1.** GO enrichment analysis of proteins interactors of a select set of SARS-CoV-2 proteins (molecular functions) (.pdf).

**Supplementary Figure S2.** GO enrichment analysis of proteins interactors of a select set of SARS-CoV-2 proteins (biological processes) (.pdf).

**Supplementary Figure S3.** Pie charts of the GO enrichment analysis of MCL clusters and clustering statistical significance of GO terms according to GoNet (cellular components) (.pdf).

**Supplementary Figure S4.** Pie charts of the GO enrichment analysis of MCL clusters and clustering statistical significance of GO terms according to GoNet (molecular functions) (.pdf).

**Supplementary Figure S5.** (.pdf).

**Supplementary Table S1.** Human PPIs from the largest connected component of the STRING-augmented network. (.xlsx)

**Supplementary Table S2.** Amino acid mapping to degenerate characters. (.xlsx)

**Supplementary Table S3.** Protein clusters of size 3 or more detected by the MCL algorithm in the STRING-augmented PPI network. (.xlsx)

**Supplementary Table S4.** Complete results from the GoNet analysis on the STRING-augmented PPI network (.xlsx).

**Supplementary Table S5.** Motifs detected as significantly clustered (FDR< 0.05) by LESMoN-Pro (.xlsx).

**Supplementary File S1.** Complete set of results of the Ontologizer GO enrichment analysis performed on the viral-host PPI network (.zip file of .txt files).

**Supplementary File S2.** Complete set of results of the MEME motif enrichment analysis performed on the viral-host PPI network (.zip file of .html files).

**Supplementary File S3.** Complete set of results of the Ontologizer GO enrichment analysis performed on the MCL clusters of the STRING-augmented PPI network (.zip file of .txt files).

**Supplementary File S4.** Complete set of results of the MEME motif enrichment analysis performed on the MCL clusters of the STRING-augmented PPI network (.zip file of .html files).

## AUTHOR INFORMATION

### Author Contributions

The manuscript was written through contributions of all authors. All authors have given approval to the final version of the manuscript. ‡These authors contributed equally.

## ACKNOWLEDGMENTS

The authors acknowledge funding from the following sources: Natural Sciences and Engineering Research Council (NSERC) of Canada Discovery grants to M.L.A. R.N. held a NSERC Canada Graduate Scholarship and an Ontario Graduate Scholarship. S. S. F. held a Canadian Institutes of Health Research Canada Graduate Scholarship. E. R. and D. N. were funded by a stipend from the NSERC CREATE in Metabolomics Advanced Training and International Exchange (MATRIX) Program. A.S. I.A. and Y.-E.C. held a NSERC Undergraduate Student Research Award. This research was enabled in part by support provided by Compute Ontario (https://computeontario.ca/) and Compute Canada (www.computecanada.ca) with Resource Allocation to M.L.A.

## ABBREVIATIONS

FDR: False Discovery Rate
GO: Gene Ontology
MCL: Markov Clustering algorithm
MTOC: Microtubule Organizing Center
PPI: Protein-Protein Interaction.

