## Supplementary Figure S1 for "Computational identification of human biological processes and protein sequence motifs putatively targeted by SARS-CoV-2 proteins using protein-protein interaction networks"

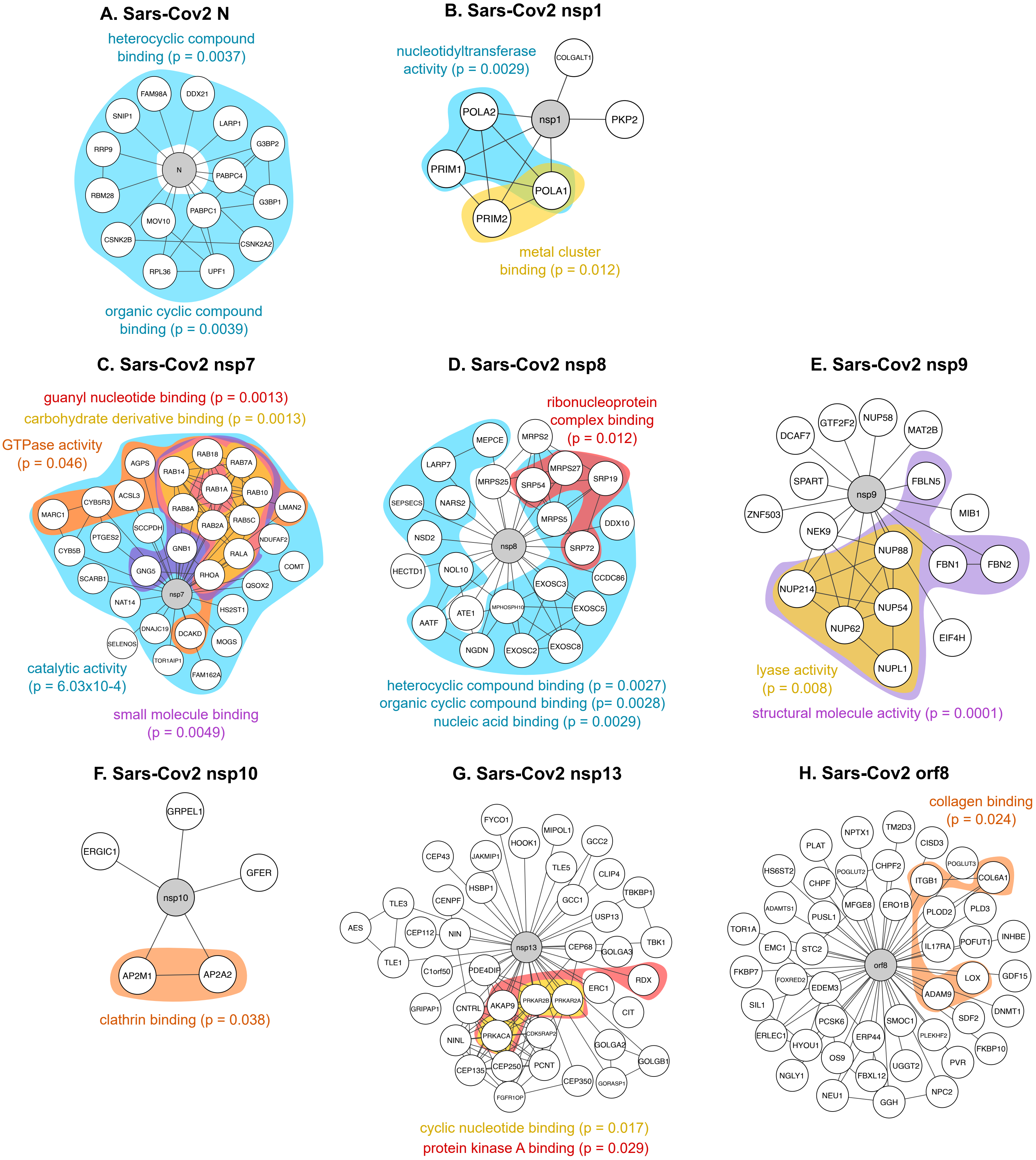


Supplementary Figure S1. GO enrichment analysis of proteins interactors of a select set of SARS-CoV-2 proteins. A) – H): GO molecular functions are color-coded. p corresponds to an FDR-adjusted p-values (See methods).
