## Supplementary Figure S3 for "Computational identification of human biological processes and protein sequence motifs putatively targeted by SARS-CoV-2 proteins using protein-protein interaction networks"

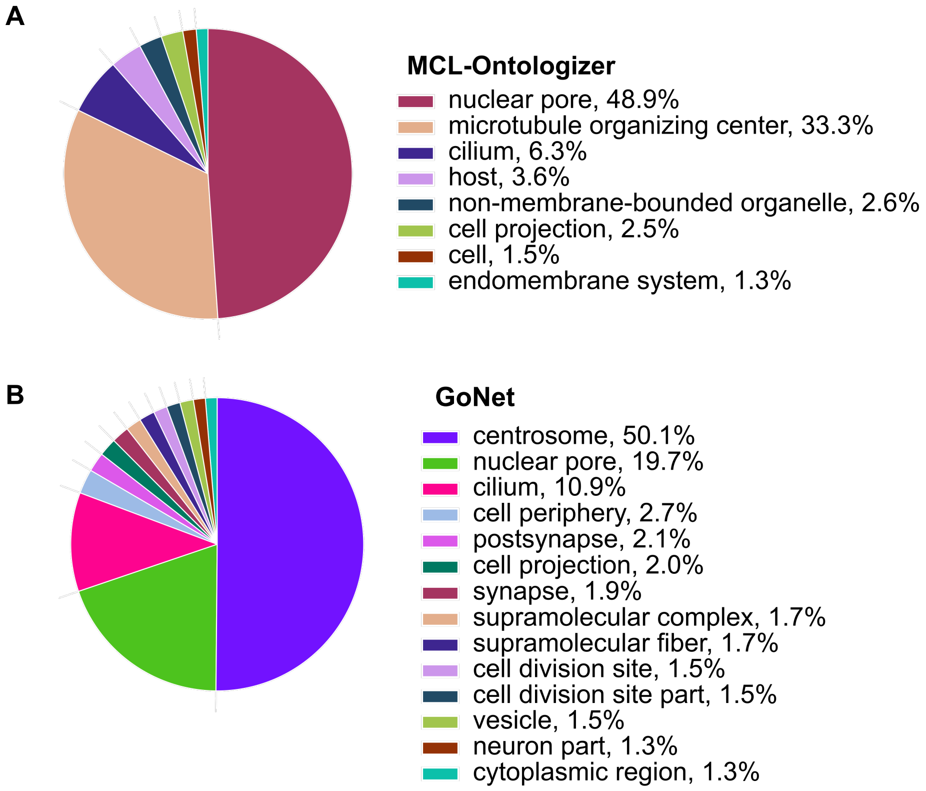


Supplementary Figure S3. Pie charts of the GO enrichment analysis of MCL clusters (A) and clustering statistical significance of GO terms according to GoNet (B). GO cellular components with the highest level of enrichment statistical significance are attributed larger pieces of the pie. The portion occupied by each GO term is also represented in percentages next to the term names.
