## Supplementary Figure S5 for "Computational identification of human biological processes and protein sequence motifs putatively targeted by SARS-CoV-2 proteins using protein-protein interaction networks"

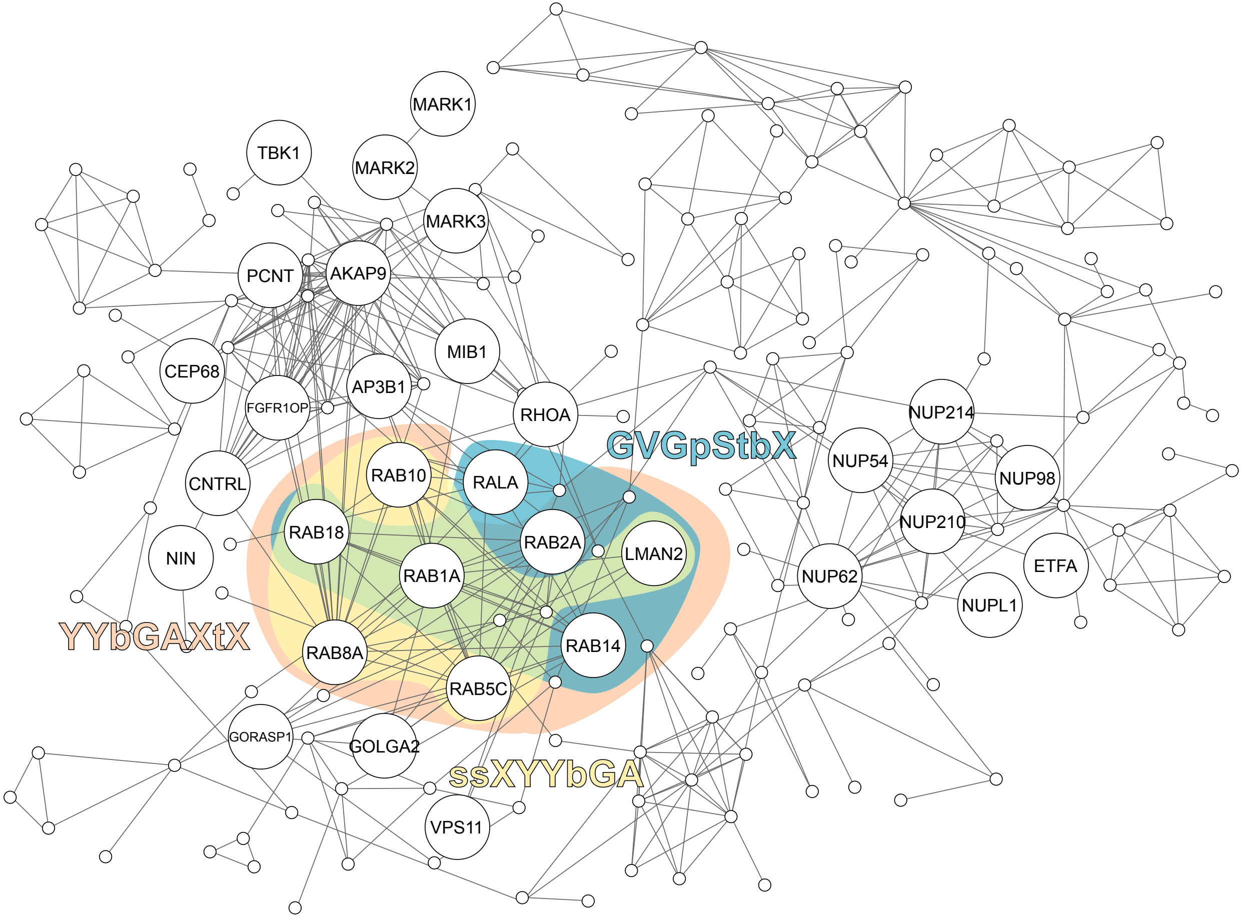


Supplementary Figure S5. Family representative sequence motifs mainly involving RAB proteins for which the associated proteins are significantly clustered in the STRING-augmented network. The complete STRING-augmented network is represented. Proteins containing significantly clustered motifs are larger and labeled. A selected set of representative motifs involving RAB proteins are shown on the network coloring the proteins containing them (FDR < 0.05).
