## Supplementary File S4 for "Computational identification of human biological processes and protein sequence motifs putatively targeted by SARS-CoV-2 proteins using protein-protein interaction networks": c20.html

MEME


MEME results in plain text format.

[
close ]

MEME results in XML format.

[
close ]

Help poup.

[
close ]

The statistical significance of the motif. MEME usually finds the most
statistically significant (low E-value) motifs first. It is unusual to
consider a motif with an E-value larger than 0.05 significant so, as an
additional indicator, MEME displays these partially transparent.

The E-value of a motif is based on its log likelihood ratio, width,
sites, the background letter frequencies (given in the command line
summary), and the size of the training set.

The E-value is an estimate of the expected number of motifs with the
given log likelihood ratio (or higher), and with the same width and site
count, that one would find in a similarly sized set of random
sequences (sequences where each position is independent and letters are
chosen according to the background letter frequencies).

[
close ]

The number of sites contributing to the construction of the motif.

[
close ]

The width of the motif. Each motif describes a pattern of a fixed
width, as no gaps are allowed in MEME motifs.

[
close ]

Click on the blue symbol below to reveal more information about this motif.

[
close ]

Click on the blue symbol below to reveal options allowing you
to submit this motif to another MEME Suite motif analysis program, to download this
motif in various text formats, or to download a sequence "logo" of
this motif PNG or EPS format.

###### Supported Programs

Tomtom
:   Tomtom is a tool for searching for similar known motifs.
    [manual]

MAST
:   MAST is a tool for searching biological sequence databases for
    sequences that contain one or more of a group of known motifs.
    [manual]

FIMO
:   FIMO is a tool for searching biological sequence databases for
    sequences that contain one or more known motifs.
    [manual]

GOMo
:   GOMo is a tool for identifying possible roles (Gene Ontology
    terms) for DNA binding motifs.
    [manual]

SpaMo
:   SpaMo is a tool for inferring possible transcription factor
    complexes by finding motifs with enriched spacings.
    [manual]

[
close ]

The log likelihood ratio of the motif.The log likelihood ratio is the
logarithm of the ratio of the probability of the occurrences of the motif
given the motif model (likelihood given the motif) versus their
probability given the background model (likelihood given the null model).
(Normally the background model is a 0-order Markov model using the
background letter frequencies, but higher order Markov models may be
specified via the -bfile option to MEME.).

[
close ]

The information content of the motif in bits. It is equal to the sum
of the uncorrected information content, R(), in the columns of the motif.
This is equal relative entropy of the motif relative to a uniform
background frequency model.

[
close ]

The relative entropy of the motif.

re = llr / (sites \* ln(2))

[
close ]

The Bayes Threshold.

[
close ]

The strand used for the motif site.

+
:   The motif site was found in the sequence as it was supplied.

-
:   The motif site was found in the reverse complement of the supplied sequence.

[
close ]

The position in the sequence where the motif site starts. If a motif
started right at the beginning of a sequence it would be described as
starting at position 1.

[
close ]

The probability that an equal or better site would be found in a
random sequence of the same length conforming to the background letter
frequencies.

[
close ]

A motif site with the 10 flanking letters on either side.

When the site is not on the given strand then the site
and both flanks are reverse complemented so they align.

[
close ]

The name of the sequences as given in the FASTA file.

The number to the left of the sequence name is the position
of the sequence in the input sequence file.

[
close ]

These are the motif sites predicted by MEME and used to build the motif.

These sites are shown in solid color and hovering the cursor
over a site will reveal details about the site. Only sequences
that contain a motif site are shown.

[
close ]

These are the motif sites predicted by MEME plus
any additional sites detected using a motif scanning
algorithm.

These MEME sites are shown in solid color and
additional scanned sites are shown in transparent color.
Hovering the cursor over a site will reveal details about the site.
Only sequences containing a predicted or scanned motif site are shown.

The scanned sites are predicted using a
log-odds scoring matrix constructed from the MEME sites.
Only scanned sites with position *p*-values less
than 0.0001 are shown.

[
close ]

These are the same sites as shown by selecting the
"Motif Sites + Scanned Sites" button except that all
sequences, including those with no sites, are included
in the diagram.

[
close ]

This is the combined match *p*-value.

The combined match *p*-value is defined as the probability that a
random sequence (with the same length and conforming to the background)
would have position *p*-values such that the product is smaller
or equal to the value calculated for the sequence under test.

The position *p*-value is defined as the probability that a
random sequence (with the same length and conforming to the background)
would have a match to the motif under test with a score greater or equal
to the largest found in the sequence under test.

Hovering your mouse over a motif site in the motif location
block diagram will show its position *p*-value and other information
about the site.

[
close ]

Use this button to download the "Motif Locations" block diagrams
as a PDF image suitable for publication.

Only the block diagrams currently visible in the inner scrolling
window (below) will be included in the image, and the numbers to
the left of each sequence name will not be included in the image.
You can change the size of the inner scrolling by moving the bottom
of the main document window up and down. You can display more
diagrams by making your browser's font size smaller.

[
close ]

Use this button to download the "Motif Locations" block diagrams
as a SVG image use in HTML documents.

Only the block diagrams currently visible in the inner scrolling
window (below) will be included in the image, and the numbers to
the left of each sequence name will not be included in the image.
You can change the size of the inner scrolling by moving the bottom
of the main document window up and down. You can display more
diagrams by making your browser's font size smaller.

[
close ]

This button will only function if your browser was
connected to the internet when you loaded this page.

To use this button, make sure your browser is connected to the internet
and then reload this page. (You may need to do a "hard refresh" to clear the cache.
On Mac, hold down the Shift key and click the Reload button.
On Windows/Linux, hold down Ctrl and press F5.)

[
close ]

This diagram shows the location of motif sites.

Each block shows the position and strength of a motif
site. The height of a block gives an indication of the
significance of the site as taller blocks are more significant.
The height is calculated to be proportional to the negative
logarithm of the *p*-value of the site, truncated at
the height for a *p*-value of 1e-10.

For complementable alphabets (like DNA), sites on the
positive strand are shown above the line,
sites on the negative strand are shown below.

Placing the cursor
over a motif site will reveal more information about the site
including its position *p*-value. (See the help
for the *p*-value column for an explanation of position
*p*-values.)

[
close ]

The name of the file(s) of sequences input to MEME.

[
close ]

The position specific priors file used by MEME to find the motifs.

[
close ]

The alphabet used by the sequences.

[
close ]

The number of FASTA sequences provided in this input file.

[
close ]

The number of characters in the sequences provided in this FASTA input file.

[
close ]

The name of the alphabet symbol.

[
close ]

The frequency of the alphabet symbol in the dataset.

[
close ]

The frequency of the alphabet symbol as defined by the background model.

[
close ]

|  |  |
| --- | --- |
| Motif | 1 |
| *p*-value | 8.23e-7 |
| Start | 23 |
| End | 33 |

###### Scanned Site

|  |  |
| --- | --- |
| Motif | 1 |
| *p*-value | 8.23e-7 |
| Start | 23 |
| End | 33 |

.

↥

⇢

*E*-value:

Site Count:

Width:

StandardReverse
Complement

Log Likelihood Ratio:

Information Content:

Relative Entropy:

Bayes Threshold:

x

#### Submit or Download

⇧⬆

⇩⬇

Submit MotifDownload MotifDownload Logo

###### Submit to program

|  |  |  |
| --- | --- | --- |
|  | Tomtom | Find similar motifs in published libraries or a library you supply. |
|  | FIMO | Find motif occurrences in sequence data. |
|  | MAST | Rank sequences by affinity to groups of motifs. |
|  | GOMo | Identify possible roles (Gene Ontology terms) for motifs. |
|  | SpaMo | Find other motifs that are enriched at specific close spacings which might imply the existence of a complex. |

Format:

Count Matrix
Probability Matrix
Minimal MEME
FASTA
Raw

|  |  |
| --- | --- |
| Format: | PNG (for web) EPS (for publication) |
| Orientation: | Normal Reverse Complement |
| Small Sample Correction: | Off On |
| Width: | cm |
| Height: | cm |

### MEME

#### Multiple Em for Motif Elicitation

For further information on how to interpret these results please access
http://meme-suite.org.   
To get a copy of the MEME software please access
http://meme-suite.org.

If you use MEME in your research, please cite the following paper:  

Timothy L. Bailey and Charles Elkan,
"Fitting a mixture model by expectation maximization to discover motifs in biopolymers",
*Proceedings of the Second International Conference on Intelligent Systems
for Molecular Biology*, pp. 28-36, AAAI Press, Menlo Park, California, 1994.
[pdf]

Discovered Motifs
  |  
Motif Locations
  |  
Inputs & Settings
  |  
Program Information
  |  
Results in Text Format  
  |  
Results in XML Format


### Javascript is required to view these results!

### Your browser does not support canvas!

#### Discovered Motifs

Please wait... Loading...

If the page has fully loaded and this message does not disappear then an error may have occurred.

#### Motif Locations

Please wait... Loading...

If the page has fully loaded and this message does not disappear then an error may have occurred.

#### Inputs & Settings

###### Sequences

| Role | Source | PSP Source | Alphabet | Sequence Count | Total Size |
| --- | --- | --- | --- | --- | --- |
| Primary Sequences |  |  |  |  |  |
| Control Sequences |  |  |  |  |  |

###### Background Model


###### Other Settings

|  |  |
| --- | --- |
| Motif Site Distribution | ZOOPS: Zero or one site per sequence OOPS: Exactly one site per sequence ANR: Any number of sites per sequence |
| Objective Function |  |
| Starting Point Function |  |
| Site Strand Handling | This alphabet only has one strand Sites must be on the given strand Sites may be on either strand |
| Maximum Number of Motifs |  |
| Motif E-value Threshold |  |
| Minimum Motif Width |  |
| Maximum Motif Width |  |
| Minimum Sites per Motif |  |
| Maximum Sites per Motif |  |
| Bias on Number of Sites |  |
| Sequence Prior | Simple Dirichlet Dirichlet Mixture Mega-weight Dirichlet Mixture Mega-weight Dirichlet Mixture Plus Add One |
| Sequence Prior Source |  |
| Sequence Prior Strength |  |
| EM Starting Point Source | From substrings in input sequences From strings on command line (-cons) |
| EM Starting Point Map Type | Uniform Point Accepted Mutation |
| EM Starting Point Fuzz |  |
| EM Maximum Iterations |  |
| EM Improvement Threshold |  |
| Maximum Search Size |  |
| Maximum Number of Sites for E-values |  |
| Trim Gap Open Cost |  |
| Trim Gap Extend Cost |  |
| End Gap Treatment | No cost Same cost as other gaps |
| Show Advanced Settings Hide Advanced Settings | |

###### MEME version

(Release date: )

###### Command line
